# Vulcan: Improved long-read mapping and structural variant calling via dual-mode alignment

**DOI:** 10.1101/2021.05.29.446291

**Authors:** Yilei Fu, Medhat Mahmoud, Viginesh Vaibhav Muraliraman, Fritz J. Sedlazeck, Todd J. Treangen

**Affiliations:** Department of Computer Science, Rice University, Houston, TX 77005, USA; Human Genome Sequencing Center, Baylor College of Medicine, Houston, TX 77030, United States of America; Department of Molecular and Human Genetics, Baylor College of Medicine, Houston, TX 77030, United States of America

**Keywords:** Long reads, Read mapping, Gap penalty, Structural variation

## Abstract

**Background:** Long-read sequencing has enabled unprecedented surveys of structural variation across the entire human genome. To maximize the potential of long-read sequencing in this context, novel mapping methods have emerged that have primarily focused on either speed or accuracy. Various heuristics and scoring schemas have been implemented in widely used read mappers (minimap2 and NGMLR) to optimize for speed or accuracy, which have variable performance across different genomic regions and for specific structural variants. Our hypothesis is that constraining read mapping to the use of a single gap penalty across distinct mutational hotspots reduces read alignment accuracy and impedes structural variant detection.

**Findings:** We tested our hypothesis by implementing a read mapping pipeline called Vulcan that uses two distinct gap penalty modes, which we refer to as dual-mode alignment. The high-level idea is that Vulcan leverages the computed normalized edit distance of the mapped reads via e.g. minimap2 to identify poorly aligned reads and realigns them using the more accurate yet computationally more expensive long read mapper (NGMLR). In support of our hypothesis, we show Vulcan improves the alignments for Oxford Nanopore Technology (ONT) long-reads for both simulated and real datasets. These improvements, in turn, lead to improved accuracy for structural variant calling performance on human genome datasets compared to either of the read mapping methods alone.

**Conclusions:** Vulcan is the first long-read mapping framework that combines two distinct gap penalty modes, resulting in improved structural variant recall and precision. Vulcan is open-source and available under the MIT License at https://gitlab.com/treangenlab/vulcan

## Background

The advent of long-read DNA sequencing over the past decade has led to many new insights in genomics and genetics [1–3]. One of the main advantages of long-read sequencing is for human research given the size and complexity of the human genome, and specifically for the detection of Structural Variation (SV) [1,2,4,5]. SVs are often defined as 50bp or larger genomic alterations that can be categorized into five types: deletions, duplications, insertions, inversions, and translocations [6,7]. Due to higher false positive and false negative rates in SV detection with short-reads, long-reads are preferred to accurately detect and fully resolve SVs [6].

In recent years, three types of long-reads have been established, produced by two sequencing platforms: Pacific Biosciences (PacBio) and Oxford Nanopore Technology (ONT)[3]. The latest PacBio device (Sequel II [3] not only sequences continuous long-reads (CLR) that have error rates of up to 10%, but also longer average length, it can also produce HiFi reads [8]. The latter is produced by repeatedly sequencing the same molecule multiple times (10-20kbp long) producing a consensus read that lowers the sequencing error down to 1% or even lower [8]. ONT is the other long-read sequencing platform. ONT also offers single-molecule sequencing and can produce ultra-long reads (>100 kbp and up to 2Mbp) [9] with drastically reduced cost with respect to HiFi reads but at a higher error rate (3-10%) [10]. In recent years, SVs have been shown as an important type of genomic alteration often leading to more modified base pairs than single nucleotide variants (SNVs) on their own [6,8]. Furthermore, SVs have been shown to have an impact on many human diseases and also other phenotypes across multiple species [6,11–13]. Most of the existing SV detection approaches depend on long-reads to facilitate the mapping of these reads to a known reference genome.

We define read mapping as the process of performing a pairwise alignment between a read and a reference genome to identify the region of origin for this DNA molecule [14,15]. Early on BLASR[16] was the method of choice for high-error long read mapping. Given its advantageous speed, BWA-MEM[17] later emerged as the method of choice to align single-molecule sequencing reads. We have previously shown that while BWA-MEM performs well in aligning these long-reads, it produces less optimal alignments in the presence of structural variants (SVs) [2,18]. This is mainly due to sequencing errors coupled with SV signals in repetitive regions are mixed and causes sub-optimal pairwise alignments, hindering an accurate detection of SV. To circumvent this issue we introduced NGMLR [2], which made use of a convex scoring matrix to better distinguish between read error and SV signal. Using this approach we were able to achieve high accuracy SV detection and at a similar speed compared to BWA-MEM. However, as sequencing throughput increased, NGMLR was not fast enough to keep up with the sheer volume of data, thus becoming a bottleneck in the analysis of larger data sets. Minimap2 [18] has since emerged as a highly-efficient long-read mapper, implementing a much faster alignment approach with extending the traditional affine gap cost model to a 2-piece affine gap model [19] and implementing an efficient chaining process. Thanks to these important algorithmic enhancements, minimap2 achieved a faster run time at a similar accuracy to state-of-the-art long-read mappers [18]. There exist several other long-read aligners that have prioritized accuracy, sensitivity, or speed, such as MashMap [20], LAST [21], GraphMap [22], and LRA[23]. However, despite promising recent progress exemplified by these methods, there is still room for improvement in long-read mapping [14].

We posit that a single strategy may not be sufficient for those variable regions; we explore in this study whether distinct heuristics implemented in the different mappers perform better or worse in certain organisms or even regions of the genome (e.g. human). The latter is especially relevant if one considers the different mutational rates per specific genomic region due to recombination [24], housekeeping genes [25], and orphan genes [26]. For example, a conserved housekeeping gene will have a very different mutational landscape compared to genes involved in immune responses (e.g. HLA[27], KIR, etc.) or compared to other highly variable genes among the human population (e.g. LPA[28], Cyp2d6).

To cope with these challenges, in this work we describe a unified long read mapping framework called Vulcan that melds alignment strategies from two different long-read mappers, here NGMLR and minimap2. At its core, Vulcan is based on the following straightforward idea: *use distinct gap penalties for different mappings between long reads and a reference genome*. Notably, Vulcan is the first long-read mapping framework that combines two scoring functions, as shown in **Figure 1**. Vulcan first maps reads starting with the fastest long-read mapper (minimap2 by default). The key idea behind Vulcan is to identify reads that are sub-optimally aligned based on edit distance and then realign them with a more sensitive (NGMLR by default). Previous works have shown that edit distance based approaches may have an effect on effective detection of SVs [26,29–31]. Here we show that edit distance can be used as a prior for sub-optimally aligned reads, highlighting the utility and accuracy of Vulcan based on NGMLR and minimap2. We apply Vulcan on simulated and real data sets (HG002) to measure the improvements of our dual-mode alignment approach in both the number of correctly aligned reads and run time. Furthermore, to showcase the benefit of improved read mappings, we compared SV calling from Vulcan mapped reads to both NGMLR and minimap2 mapped reads on simulated ONT reads and Human ONT, Human PacBio CLR and HiFi reads.

**Fig 1.**
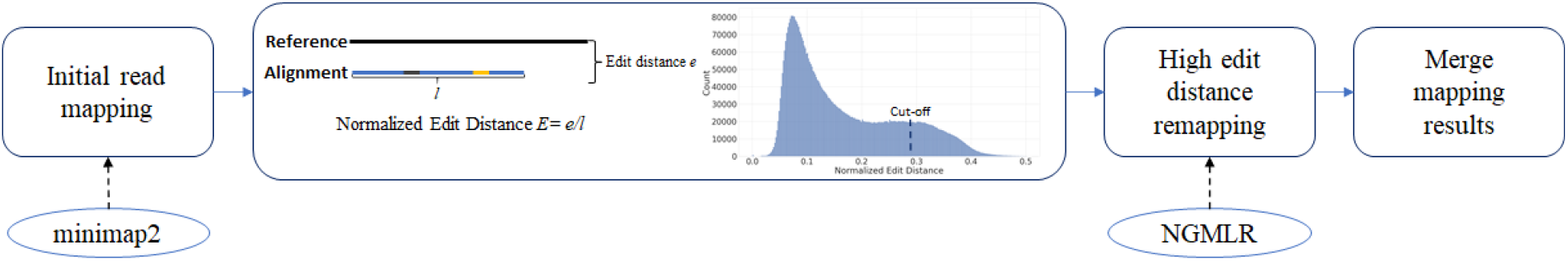
Overview of Vulcan: As step 1, Vulcan takes raw ONT or Pacbio reads as input, then uses minimap2 to map them to the provided reference genome. Subsequently, in step 2, Vulcan performs a normalized edit distance calculation to identify the reads with the highest normalized edit distances. In step 3, Vulcan realigns the high edit distance reads with NGMLR. Finally, in step 4 Vulcan merges the minimap2 and NGMLR remapped reads to create a new bam file.

## Data Description

To evaluate Vulcan’s ability to improve structural variant calling, we simulated five types of structural variant in the reference genome (*Saccharomyces cerevisiae* S288C). Specifically, we selected *Saccharomyces cerevisiae* S288C genome as the reference, and added structural variations into the genome with SURVIVOR(1.0.7), simSV [11], later, we used Nanosim-h (1.1.0.4) [32] to simulate 10X coverage reads set. We ran NGMLR, minimap2 and Vulcan on the dataset and used Sniffles (version 1.12) to identify SV. In this experiment we also included other SV types like duplication (DUP), translocation (TRA) and inversion (INV).

Additionally, we used real data to show the improvements over HG002, a benchmark sample well studied by GIAB (NIST). Here we downloaded ONT, PacBio HiFi and PacBio CLR data sets for the same sample. The data is available at https://ftp.ncbi.nih.gov/giab/ftp/data/AshkenazimTrio/HG002_NA24385_son/, and has been described in multiple papers[33,34]. The subsample of coverages was performed with seqtk[35].

## Analyses

To demonstrate the ability of Vulcan to improve the overall mapping of long-reads and thus to improve the structural variant detection across organisms we used simulated (*S. cerevisiae S288C*) and real data (human hg19) datasets. For the real datasets we utilized three distinct long-read technologies (PacBio: HiFi and CLR, Oxford Nanopore) [32,34]. Using these datasets, we evaluated the edit distance improvement after Vulcan’s refinement and SV calling performance (recall, precision and F1 score). Also, we show that Vulcan reduces computational time against the methods that use convex gap penalty (NGMLR).

### Vulcan Improves Long-read Mapping over Minimizing Read-to-Reference Edit Distance

First, we investigated Vulcan’s ability to identify reads that would benefit from convex gap penalty vs two-piece affine gap penalty by thresholding the reported edit distance from the mappers (see Methods section), and thus minimize the edit distance between the read and mapped location on the reference genome. To accomplish this, we mapped the GIAB HG002 ONT Ultra-long UCSC dataset using minimap2 and investigated the alignments from the reads given their reported edit distance (NM tag).

We benchmarked Vulcan genome wide to see if it improves the overall edit distance compared to minimap2 alone. Figure 2A shows this trend as Vulcan on the median has a lower normalized edit distance than minimap2 alone. Notably, Vulcan does not recapitulate the overall distribution of edit distance from NGMLR as it only realigns 10% of the reads in this example. Thus, by automatically realigning only 10% of the reads Vulcan significantly improves the alignments in certain regions of the human genome compared to minimap2. These results provide support for our dual-mode alignment strategy implemented in Vulcan to select reads based on their normalized edit distance and then realigning these using NGMLR seem to work and indeed improve the representation of SV (Table 1 and 2).

**Table 1.**
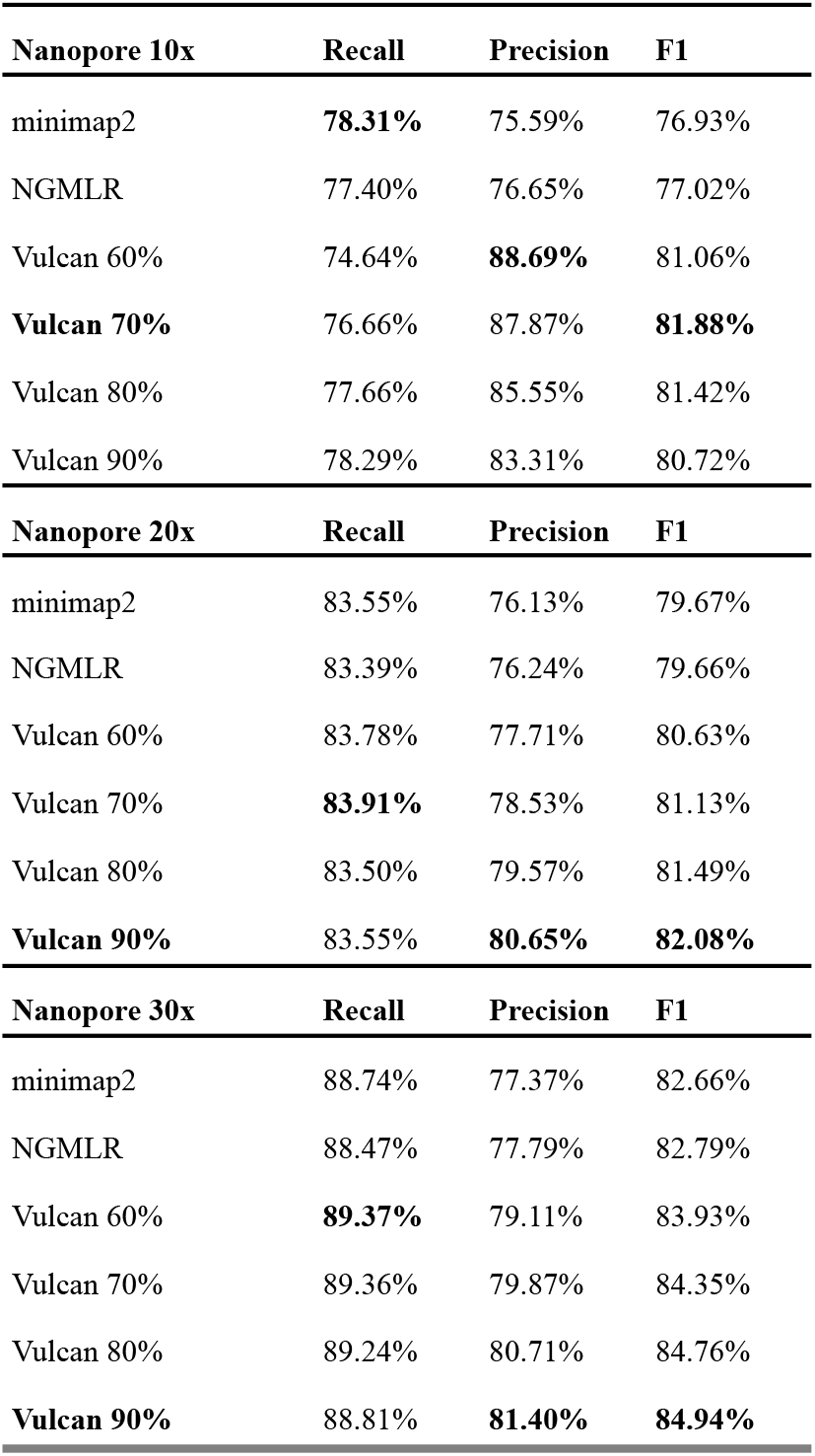
Benchmarking SV recall, precision, and F1 on HG002 Human (hg19) ONT reads at varying coverages (10X, 20x, 30x). Various percentile cut-offs for Vulcan were used, including: 60%, 70%, 80%, 90%. SV calls based on Vulcan mappings achieve the highest F1 score for various cut-off values.

**Table 2.** Benchmarking SV recall, precision, and F1 on HG002 Human (hg19) PacBio reads (CLR and HiFi) at varying coverages (CLR 10x, 20x, 30x; HiFi 10x). Various percentile cut-offs for Vulcan were used, including: 60%, 70%, 80%, 90%. Vulcan achieves the highest F1 score on PacBio CLR 20X and 30X reads, with minimap2 achieving the highest F1 score on PacBio CLR 10X and PacBio HiFi 10X reads.

| PacBio CLR 10x | Recall | Precision | F1 |
| --- | --- | --- | --- |
| **minimap2** | **62.85%** | 88.88% | **73.63%** |
| NGMLR | 60.11% | 86.44% | 70.91% |
| Vulcan 60% | 60.13% | 89.41% | 71.90% |
| Vulcan 70% | 60.79% | 90.12% | 72.61% |
| Vulcan 80% | 60.97% | **90.13%** | 72.73% |
| Vulcan 90% | 61.85% | 89.93% | 73.29% |
| PacBio CLR 20x | Recall | Precision | F1 |
| minimap2 | **77.76%** | 71.85% | 74.69% |
| NGMLR | 75.74% | 68.36% | 71.86% |
| Vulcan 60% | 75.74% | 74.69% | 75.21% |
| Vulcan 70% | 75.98% | 75.32% | 75.65% |
| Vulcan 80% | 76.22% | **75.65%** | 75.93% |
| **Vulcan 90%** | 76.90% | 75.08% | **75.98%** |
| PacBio CLR 30x | Recall | Precision | F1 |
| minimap2 | **83.71%** | 86.25% | 84.96% |
| NGMLR | 81.79% | 82.41% | 82.10% |
| Vulcan 60% | 82.05% | 86.31% | 84.12% |
| Vulcan 70% | 82.33% | 87.12% | 84.66% |
| Vulcan 80% | 82.47% | **87.60%** | 84.96% |
| **Vulcan 90%** | 82.75% | 87.49% | **85.05%** |
| PacBio HiFi 10X | Recall | Precision | F1 |
| **minimap2** | **81.50%** | **90.70%** | **85.85%** |
| NGMLR | 78.22% | 86.26% | 82.04% |
| Vulcan 60% | 77.73% | 86.04% | 81.68% |
| Vulcan 70% | 77.74% | 86.19% | 81.75% |
| Vulcan 80% | 76.40% | 85.75% | 80.81% |
| Vulcan 90% | 76.26% | 85.73% | 80.72% |

**Fig. 2.**
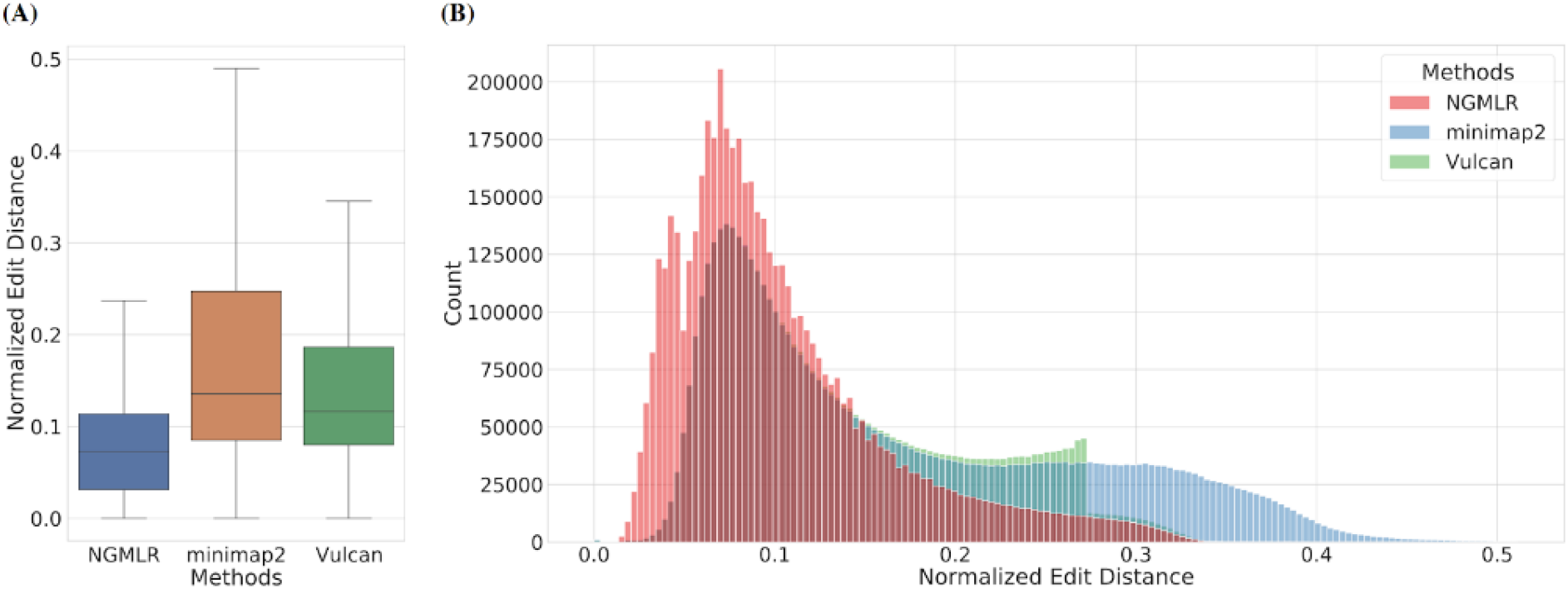
Overall edit distance improvements. **A:** Normalized edit distance comparison of Vulcan’s 90% percentile cut-off, NGMLR and minimap2’s mapping result with human ONT 30X reads. We can see clear evidence that the realignment of only 10% of the reads lead to an improvement in edit distance and thus of the variant calling. **B:** Distribution of mappings’ normalized edit distance from Vulcan, NGMLR and minimap2. Vulcan has a lower edit distance mapping than minimap2 with NGMLR’s refinement.

### Vulcan Accelerates Long-read Mapping for SV Calling

Next, we evaluated the speedup of Vulcan compared to minimap2 and NGMLR. As shown in Figure 3, Vulcan is able to achieve up to a 4X speedup over NGMLR, from 778 CPU hours down to 191 CPU hours for the 90% cut-off (default parameter setting for human genome mapping). When increasing the edit distance cut-off percentile, Vulcan runtime decreases linearly. When comparing to minimap2’s runtime we see that Vulcan’s default setting only requires 2.5X more CPU hours compared to over 10X more CPU hours required for NGMLR. This highlights Vulcan’s ability to drastically reduce NGMLR runtime and maintain comparable runtime to minimap2, one of the most efficient long-read aligners that currently exists. The RAM usage of Vulcan 90% cut-off with PacBio CLR 20X is 42.652 GB.

**Fig. 3.**
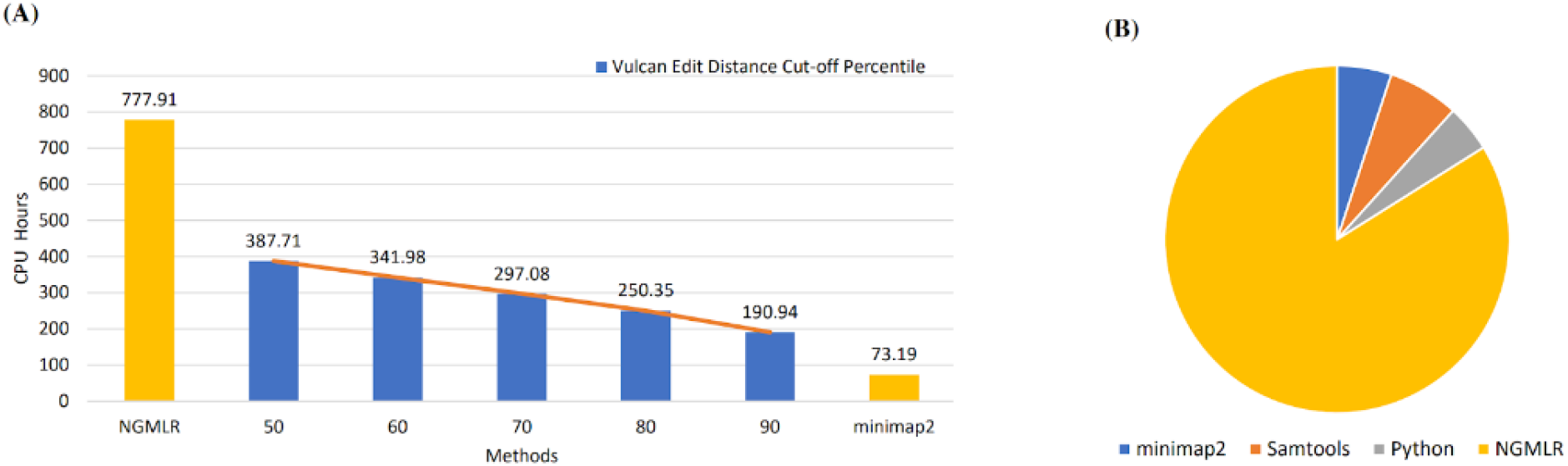
Comparing runtime for Vulcan, NGMLR and minimap2. Runtime was measured in CPU hours for all programs. **A:** Vulcan achieves an approximately linear acceleration with the increase of the cut-off percentile. With 90% percent cut-off, Vulcan only takes less than 1/4 time that NGMLR uses. **B:** The majority of Vulcan’s runtime is spent in running NGMLR on the subset of reads leading to an improvement of their alignments.

We also show the relative contribution to CPU hours for each component in Vulcan (Figure 3B): minimap2, samtools, file parsing and edit distance calculation with Python, and NGMLR. As expected, NGMLR dominates this breakdown when mapping the reads that are above the Vulcan cut-off (90% in this experiment), the remaining components represent minor contributions to Vulcan’s execution time.

### Structural Variation Detection Benchmarking

Next, we highlight NGMLR’s SV-aware mappings enable the improved detection of SV (here deletion indicated by black lines in IGV) compared to the mapping results from minimap2 (Figure 4A). We see in this example that minimap2 demonstrates a more scattered pattern of the deletion signal across all three regions (Figure 4A, 4B, and 4C). These regions include an insertion and a deletion, which induce noisy alignments from minimap2. In contrast, automatically realigning the reads with Vulcan using NGMLR shows a more consistent mapping pattern (Figure 4B and 4C). Notably, Vulcan is able to eliminate a false positive SV call by preferentially selecting a convex gap penalty over the two-piece affine gap penalty (Figure 4C), highlighting the benefit of trading off increased runtime (measured in CPU hours) for increased accuracy (measured as fewer SV false positives).

**Fig. 4:**
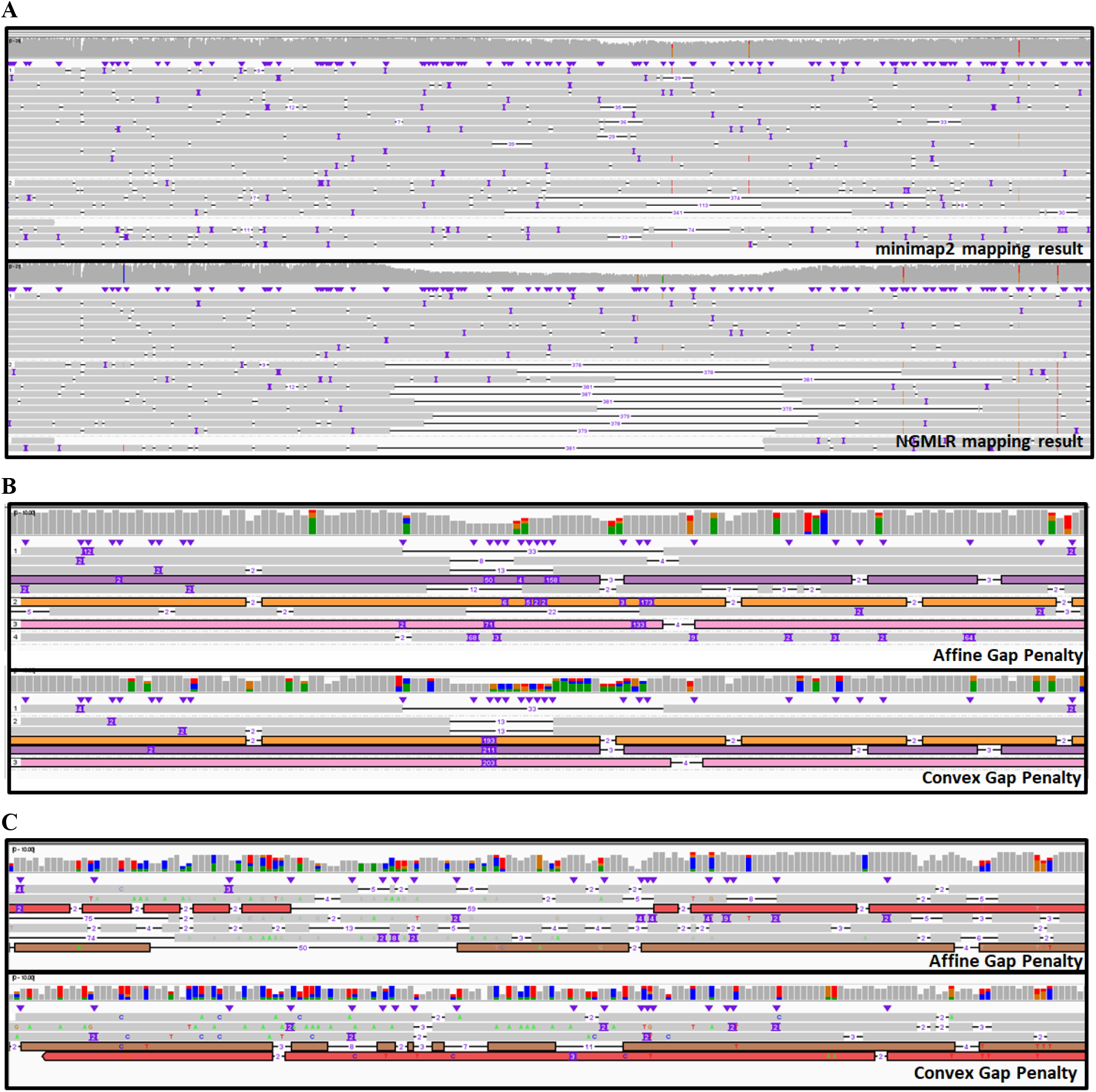
Comparison of the two read mappers used in Vulcan based on 30x ONT data. **A:** An example at chr2:112,870,823-112,871,894 of reads that show a higher normalized edit distance and thus were automatically realigned with NGMLR. The overall alignments of these reads improved, clearly highlighting a larger deletion at this location compared to the minimap2 alignments. **B:** Another example at chr1:108,567,498-108,567,633 of automatically aligned reads with Vulcan. The colored reads indicated the same read aligned by the two different methods. The realignment with NGMLR clearly shows a deletion and insertion to be present likely on the two different haplotypes. **C:** Example false positive SV call improved by Vulcan mapping. This is an example of a false positive SV call based on minimap2 that would later be resolved with Vulcan’s alignment. The region of the genome is on chr1 at 167,978,740.

### Benchmarking Structural Variant calling with Vulcan’s mappings on simulated ONT data

To follow up on the previous result, we next benchmarked SV calls based on each of the three mapping strategies: minimap2, NGMLR, and Vulcan. To perform this evaluation, we simulated Nanopore reads from the *Saccharomyces cerevisiae S288C* genome. As we see in Figure 5, Vulcan combined with Sniffles offers the highest recall and lowest FDR of all three mapping approaches. Next, Figure 5B highlights that Vulcan has the highest recall for all five SV types. We see that minimap2 has the lowest recall for duplications on this low coverage simulated long read dataset. However, both NGMLR and Vulcan are able to capture the duplication with greater than 99% recall. We also see that while translocation and insertion SV recall is identical for all three mapping approaches, Vulcan mappings help to improve both inversion and deletion detection. With respect to precision (Figure 5C), Vulcan once again performs best across all five SV categories, with NGMLR mirroring Vulcan performance in all cases.

**Fig. 5.**
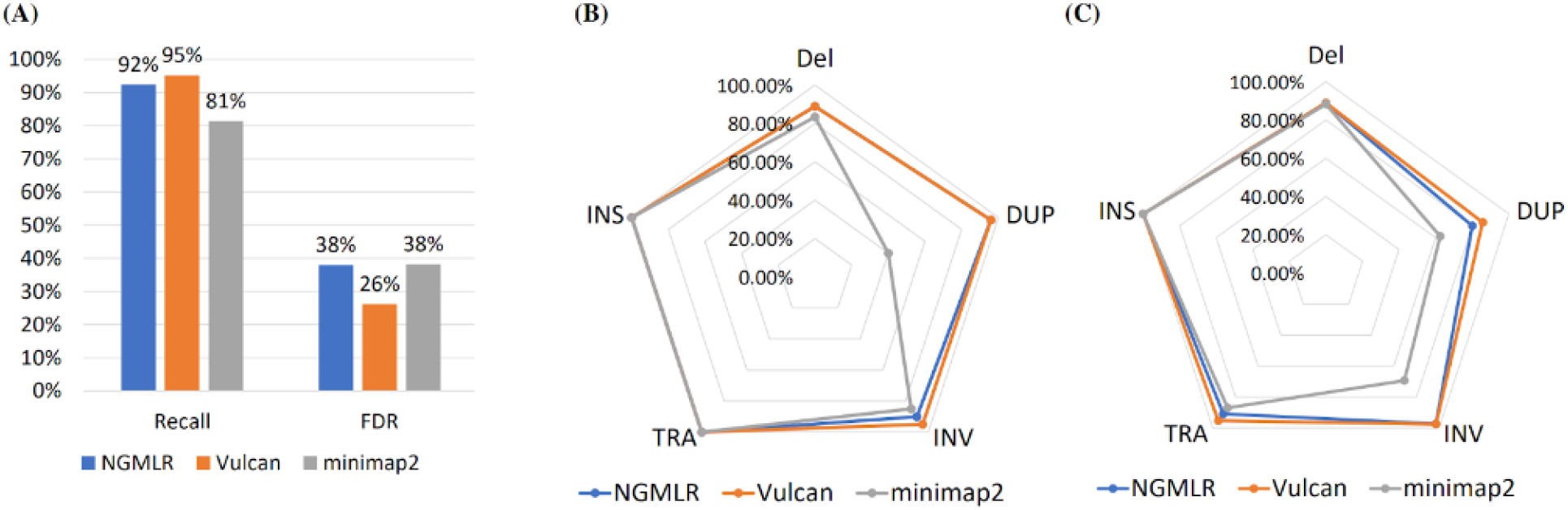
Benchmarking SV calls on simulated structural variants (INS: insertions, Del: deletions, TRA: translocations, DUP: duplications and INV: inversions) with ONT reads simulated from Saccharomyces Cerevisiae. **A:** Recall and false discovery rate (FDR) of Sniffles’ SV calling on simulated Nanopore reads with three different mappers. SV calls on Vulcan mappings offer the highest recall (95%) and lowest FDR (26%). **B:** Recall of different SV types from minimap2, NGMLR, and Vulcan mappings with Sniffles’ SV calling on simulated Nanopore reads. **C:** Precision of different SV types from NGMLR, Vulcan and minimap2’s mappings with Sniffles’ SV calling on simulated Nanopore reads. NGMLR has similar performance across all SV types, while minimap2 has a lower precision on inversions and duplications.

### Benchmarking Structural Variant calling with Vulcan’s mappings on GIAB human data

Given the promising SV calling results based on Vulcan mappings we have discovered in the simulated data, we next evaluated SV calling using Vulcan on real Human (hg19) read samples from the Genome In A Bottle (GIAB) project [34]. Similar to the SV experiment with simulated data, we used Sniffles to call SVs called from Human (hg19) reads mapped from each of the three methods: minimap2, NGMLR, and Vulcan. This GIAB dataset allows us to evaluate against an established ground truth on real hg19 long-read sequencing data. We will next describe SV performance for various Nanopore coverages (20x, 30x, 50X), PacBio CLR (20x), and PacBio HiFi (10X) datasets.

Specifically, we tested Vulcan on three different coverages across ONT and PacBio data sets with respect to improving the SV calling ability based on the GIAB SV call sets. **Table 1** shows the performance for Vulcan, NGMLR and minimap2 together with Sniffles to identify SV across the data set. Similar to the simulated data, we achieve the best F1 scores using Vulcan together with Sniffles. Vulcan provides the most improvement on lower coverage data sets. For the Nanopore 20X coverage, which is roughly equivalent to one ONT Flow cell of a human genome, Vulcan improves F1 score by 3.13% compared to minimap2 based alignments.

We then benchmarked the impact of the normalized edit distance thresholds for the ONT 30x data set (**Table 1**). We show that by increasing the cut-off percentile, we realign fewer reads and thus Vulcan exhibits lower overall runtime. However, this subsequently results in lower SV recall but higher precision. We observed the highest SV recall for Vulcan with a 60% cut-off when realigning the top 40% edit distance reads. SV precision was the highest at a 90% threshold where only the top 10% of the reads are realigned. Notably, across all thresholds, Vulcan performs the best in terms of F1 score. Vulcan by default uses a 90% threshold, yielding up to a 5% improvement in F1 score on low coverage (10X) ONT data. However, SV calls based on minimap2 mappings achieved the highest recall on 10x coverage (0.02% improvement over Vulcan mappings).

Finally, we investigated Vulcan’s performance with respect to Sniffles structural variant calls on PacBio CLR and HiFi human datasets (**Table 2**). PacBio CLR and HiFi reads offer a different error profile compared to ONT reads, with PacBio HiFi representing the lowest error rate long-reads available to date. As we see in **Table 2**, structural variant calls from Vulcan mappings offer the best recall, precision, and F1 score for 20x coverage PacBio CLR data, improving on both NGMLR and minimap2 based SV calls by over 2% in F1 score and nearly a 4% improvement over minimap2 and NGMLR precision. The F1 score improvement is due to the SV calls based off of Vulcan offering similar recall to existing approaches but improved precision. However, when comparing SV recall, we see that Vulcan mappings offer slightly lower performance compared to minimap2, while meeting or exceeding NGMLR recall. We also observed that SV calls based on Vulcan mappings offer a slightly increased recall rate when the normalized edit distance cut-off increases in PacBio CLR read dataset, different from the Nanopore dataset results. One difference between these two datasets is that the coverage of the PacBio CLR dataset is lower, and so the Sniffles minimum read support is set lower. Then when increasing the cut-off percentiles for Vulcan, there remain enough NGMLR mappings to meet or exceed the minimum read number support for SV calling.

## Discussion

In this paper we have introduced Vulcan, a novel long-read mapping tool, that leverages dual-mode long-read alignment that we have shown improves SV calling. Vulcan uses the edit distance information across the mapped reads to rapidly identify regions that are better suited for a convex gap penalty vs two-piece affine gap penalty. The key idea behind Vulcan is that different regions of the genome can benefit from distinct alignment methods (e.g. due to differences in mutation rate) and thus e.g. improves SV detection. The latter is often highlighted over mismapped reads, indicated by a higher per read substitution and Indels rate [2,34]. Throughout the results section we have highlighted the benefits of using a dual-mode alignment approach compared to minimap2 and NGMLR alone; Vulcan not only results in long-reads mapped at smaller edit distances, it improves the recall and precision of SV calling on ONT data.

Our results show that Vulcan runs up to 4X faster than NGMLR alone and produces lower edit distance alignments than minimap2, on both simulated and real datasets. In addition to improved alignment (Figure 3), we also show that using Vulcan improves the precision and recall of structural variant calling for both PacBio CLR and ONT data sets (**Table 1, Table 2**). Specifically on ONT, Vulcan is able to achieve up to a 5% improvement in F1 score for SV calls (harmonic mean of recall and precision) over the other two mappers minimap2 and NGMLR. This result not only highlights the benefit of dual-mode alignment, it supports our hypothesis that Vulcan can improve SV calling in human genome samples. We further speculate that Vulcan could improve SNV calling for complex regions. However, the edit distance selection of the reads would need to be adopted for this task and as the signal would not be that clear we opted out of benchmarking this. Nevertheless, SNVs detection around breakpoints or within SV will obviously be improved.

When designing Vulcan, we opted to focus on precision and computational efficiency. The NM tag is required according to SAMtools specifications and contains all the information needed to evaluate alignment quality. Future improvements to this approach may include not counting every difference on the read (i.e. edit distance), but instead only the start of each edit. The latter would count a longer deletion as 1 and not by the length of the event as in the current implementation. Therefore, misalignments that often introduce many smaller events and or substitutions would be more penalized than larger insertion or deletion. This could slightly improve the selection process of Vulcan, but will lead to longer runtimes since the entire alignment would need to be reconstructed per read. This approach would also consume the majority of the time of our parsing method and thus significantly alter the runtime. Thus, we did not implement this in the current version of Vulcan, but will continue to investigate other filtering schemas. We currently do not use MAPQ as a filtering criteria due to the fact that MAPQ reports the confidence of a read in a region (weighted distance of best vs. other potential alignments) [36]. This is indicative for repeats or other regional properties, but not for misalignments, or misrepresentation of variants. The issue with correct or wrong representation of SV is more related to the alignment score or chosen alignment algorithm rather than the region. Most of the time NGMLR will not change the location of the read compared to Minimap2 but the alignment itself. For example, Figure 3 shows the same reads in the same region but with a better variant representation. Thus, using the normalized edit distance has shown to be a robust and rapid approximation to detect these alignment artifacts.

Finally, we note that Vulcan could be used for any combination of long-read mappers that output the edit distance (NM tag) directly within sam/bam file output. Thus, allowing the inclusion of WinnowMap [37], LAST[21], LRA[23], or Duplomap [38] may further exploit our observation that variable gap costs for different read-to-reference mappings provide improved SV calling, while offering improve runtimes compared to the more computationally expensive long-read mapping approaches.

## Potential Implications

A key finding of this manuscript is that the utilization of dual-mode alignment, combining convex gap costs with two-piece affine gap costs, leads to improvements in alignment edit distance and subsequently SV identification. Notably, we see that SV calling based on minimap2 mappings has low recall for duplications, compared to near perfect recovery of duplications with NGMLR and Vulcan. Recently, Jain *et al*. [37] discuss that the minimizer selection strategy in minimap2 may lead to a degradation in repeat detection. Improved SV calling based on Vulcan’s results can be attributed to leveraging the strengths of the long-read alignment strategies found in minimap2 and NGMLR. Vulcan provides the first approach for long-read mapping able to track variable mutation rates and predominant mutation types at certain regions or SV hotspots. The straightforward idea behind Vulcan of adapting alignment gap costs to specific regions of the genome may be found useful for compensating for highly polymorphic regions such as HLA, a 14mbp of the human genome and at the center of several recent studies [24]-[26]. Vulcan takes the first step in leveraging this observation, and we anticipate other mappers for long reads to follow up on this observation. In conclusion, in this study we have shown that combining different long-read alignment strategies improves SV recall and precision of human structural variation detection and have provided a new open-source software tool (Vulcan) that encapsulates these benefits.

### Methods

The main idea behind Vulcan is that we combine the benefits of two popular long read mapping tools (here NGMLR and minimap2) for the improved structural variant (SV) calling. To accomplish this, we first map the reads (sequenced on the Oxford ONT or PacBio platforms) to a reference genome with minimap2 (2.17-r941), then identify the large edit distance alignments taken from minimap2 mapping results and flag them for realignment with NGMLR(0.2.7). As shown in Figure 1, Vulcan is composed of four main steps: (i) Initial read mapping, (ii) Normalized Edit Distance Calculation, (iii) High edit-distance remapping (iv) merging mapping results for downstream structural variant calling. The first step of the pipeline is to map all the reads against the reference using minimap2 and its preset parameters suitable for either PacBio or Oxford Nanopore long-read sequences. Subsequently, Vulcan uses the edit distance and scans the reads. The edit distance is the number of substitutions, insertions, or deletions that are different between the read and its region of the reference [37,39]. The edit distance is captured by the “NM” tag (mandatory tag in sam format) in reads mappers used by Vulcan.. We normalize the edit distance with the overall read length to obtain a ratio that represents the alignment of a given read. By dividing the edit distance by the alignment length of a read, we can normalize it to calculate the number of mismatches given an alignment length, i.e. with longer alignments, we tolerate larger edit distances. And normalized edit distance can be expressed as *E = e/l*, where *e* is the edit distance and *l* is the alignment length. We only keep all the primary mappings and gather the normalized edit distances with SAMtools and pysam [40]. Note, the secondary mappings typically have larger edit distances as they have a lower mapping quality than the primary mapping, which may lead to the increase of high edit distance mappings in the distance profile we generated. With the knowledge of all the normalized edit distance calculated from minimap2’s mapping result, we can now set a percentile cut-off in agreement with the user’s preference (90% as the default, based on experimental results). With the selected percentile cut-off, we can separate reads mapped with minimap2 into two sets: reads that are mapped below the cut-off and reads that are mapped above the cut-off. If we only use raw edit distance, bamtools [40,41] supports splitting mapped reads via specific tags. However, with normalized edit distances, we instead use pysam, a wrapped python interface for htslib [40] to calculate the normalized edit distance and split the mapping result. We then extract all the reads above the cut-off and re-map them with NGMLR. Thanks to NGMLR’s ability to accurately remap large edit distance reads, Vulcan is able to improve minimap2’s high edit distance results (in some cases) into small edit distances read mappings. Finally, we combine the mapping results, specifically, the mapped reads from minimap2 below the cut-off and the remapped reads from NGMLR, into a final merged and sorted BAM file. Vulcan was written in Python3.8 utilizing the multiprocessing module for multithreading support. All versions of software and parameters utilized in this study are provided in Supplementary Table 1.

### Computational Benchmarking

To evaluate Vulcan’s computational performance, we assessed the fold speedup versus NGMLR and also compared Vulcan to minimap2. We chose PacBio CLR reads with 20X coverage as test data. In this experiment, we tested our speed-up curve under different edit distance cut-offs in Vulcan and compared them with NGMLR and minimap2. We used the time command in Linux to record the program running time, and the time was recorded as CPU time of program runtime. All the CPU times are converted to hours. Furthermore, in order to profile the individual steps of Vulcan, we also counted the time usage per step on PacBio 30X coverage dataset with 90% percentile normalized edit distance cut-off. In the time benchmarking experiment, the read dataset size is roughly 64 GB and contains 8,642,942 reads.

### Human Read Dataset Structural Variation Detection Evaluation

We used Vulcan on three long read human genome data sets: Nanopore Ultra Long reads, PacBio HiFi reads and PacBio CLR reads [34]. We downloaded these three long read types from Genome in a Bottle project [33], and mapped them to the human reference genome hg19. Furthermore, we used Sniffles to call SVs from our mapping result, then compared with the ground truth that GIAB provided through truvari [34].

Sniffles [2] allows users to define the minimum number of reads supported for the structural variation (SV) calling; we set that parameter as 2 and then use bcftools [42] to further filter the minimum supported read number to achieve the optimal F1 score. We set the minimum read support to be the same for all three methods when the coverage and read type is the same, and the optimal F1 score was preferentially selected for both minimap2 and NGMLR.

The experiment was performed on an Intel(R) Xeon(R) Gold 5218 CPU @ 2.00GHz with 64 threads with Ubuntu 18.04 LTS.

## Declarations

## List of abbreviations

Normalized edit distance: normalized edit distance can be expressed as *E = e/l*, where *e* is the edit distance and *l* is the alignment length.
SV: structural variants
ONT: Oxford Nanopore Technologies
PacBio CLR: PacBio Continuous long read
PacBio HiFi: PacBio circular consensus sequencing
SNV: Single nucleotide variation
INS: insertions
DEL: deletions
TRA: translocations
DUP: duplications
INV: inversions

## Ethical Approval

Not applicable.

## Consent for publication

Not applicable.

## Author’s Contributions

All authors conceived the experiment(s), analyzed the results and reviewed the manuscript. YF and MM conducted the experiment(s). F.S. and T.J.T. managed the project.

## Acknowledgments

The authors want to thank Dreycey Albin for contributing critical discussion.

## Availability of source code and requirements

- Project name: Vulcan
- Project home page: https://gitlab.com/treangenlab/vulcan
- Operating system(s): Unix
- Programming language: Python
- Other requirements: Python 3.8 or higher, conda 4.10.1
- License: MIT

## Data Availability

- The Saccharomyces cerevisiae 288C reference genome for reads and SV simulation, taxonomy ID 559292 in NCBI taxonomy database, is available at: https://www.ncbi.nlm.nih.gov/Taxonomy/Browser/wwwtax.cgi?id=559292
- Simulated reads are available at: https://rice.box.com/v/vulcandatasimulation.
- The Ashkenazim Trio HG002 raw sequence data, and ground truth sets of structural variations are available at https://ftp.ncbi.nih.gov/giab/ftp/data/AshkenazimTrio/

