## supplementary table 1 for "Vulcan: Improved long-read mapping and structural variant calling via dual-mode alignment"

### Supplementary Info

##### Supplementary Table 1: Programs, program versions, and parameters used in this study.

<table>
<tr><th>Program</th><th>Version</th><th>Usage</th><th>Parameters</th></tr>
<tr><td>Vulcan</td><td>1.0.2</td><td>read mapping</td><td>vulcan [] -t threads</td></tr>
<tr><td>minimap2</td><td>2.17-r941</td><td>simulated ONT read mapping</td><td>minimap2 -x map-ont -a</td></tr>
<tr><td>minimap2</td><td>2.17-r941</td><td>Human ONT read mapping</td><td>minimap2 -x map-ont -a -z 600,200</td></tr>
<tr><td>minimap2</td><td>2.17-r941</td><td>Human PacBio HiFi read mapping</td><td>minimap2 -a -k 19 -O 5,56 -E 4,1 -B 5 -z 400,50 -r 2k –eqx –secondary=no</td></tr>
<tr><td>minimap2</td><td>2.17-r941</td><td>Human Pacbio CLR read mapping</td><td>minimap2 -x map-pb -a –eqx -L -O 5,56 -E 4,1 -B 5 –secondary=no -z 400,50 -r 2k -Y</td></tr>
<tr><td>NGMLR</td><td>0.2.7</td><td>read mapping</td><td>NGMLR -x [ont | pacbio] –bam-fix</td></tr>
<tr><td>Sniffles</td><td>1.0.12</td><td>SV calling</td><td>sniffles -s 2</td></tr>
<tr><td>bcftools</td><td>1.7</td><td>VCF filtering</td><td>bcftools view -i (INFO/SVLEN>=50|INFO/SVLEN<=-50|INFO/SVLEN=0|INFO/SVLEN=1)&(INFO/RE={read_num})</td></tr>
<tr><td>Truvari</td><td>v2.0.0-dev</td><td>SV benchmarking</td><td>truvari bench -b {GIAB_vcf} -c {sorted_gzipped_vcf} -f {reference_genome} –passonly–giabreport –pctsim=0 –multimatch –includebed {GIAB_bed} -o {benchmarking_output}</td></tr>
<tr><td>SURVIVOR</td><td>1.0.7</td><td>SV simulation</td><td>SURVIVOR simSV {reference_genome} {parameter_file} 0.01 0 {output_prefix}</td></tr>
</table>
